# Gamma and beta bursts during working memory read-out suggest roles in its volitional control

**DOI:** 10.1101/122598

**Authors:** Mikael Lundqvist, Pawel Herman, Melissa R. Warden, Scott L. Brincat, Earl K. Miller

## Abstract

Working memory (WM) activity is not as stationary or sustained as previously thought. There are brief bursts of gamma (∼55–120 Hz) and beta (∼20–35 Hz) oscillations, the former linked to stimulus information in spiking. We examine these dynamics in relation to read-out from WM, which is still not well understood. Monkeys held a sequence of two objects and had to decide if they matched a subsequent sequence. Changes in the balance of beta/gamma suggested their role in WM control. In anticipation of having to use an object for the match decision, there was an increase in spiking information about that object along with an increase in gamma and a decrease in beta. When an object was no longer needed, beta increased and gamma as well as spiking information about that object decreased. Deviations from these dynamics predicted behavioral errors. Thus, turning up or down beta could regulate gamma and the information in working memory.

## Introduction

Sustained spiking activity has been the dominant neural model of working memory (WM)^1-5^. The idea is that, once activated by a stimulus, neurons keep spiking, sustaining the representation of that stimulus. Recent closer examinations have revealed that complex dynamics underlie sustained activity, with brief, discrete narrow-band oscillatory bursts in the gamma and beta bands^6^. Gamma bursts (∼55–120 Hz) were tied to spiking carrying information about the remembered items. Beta bursts (∼20–35 Hz) were associated with suppression of both informative spiking and gamma. These data are consistent with a model in which gamma-associated spiking stores memories by short-term changes in synaptic weights^7^. Multiple items can be held in WM without mutual interference because different gamma bursts store different items.

The model also predicts that gamma bursting plays a role in WM read-out. This is difficult to test in many tasks because read-out often coincides with a behavioral response, adding a confounding or obscuring motor component. Thus, we turned to data from an experiment with a unique design^8, 9^. The animals determined if a test sequence of two objects matched an earlier sample sequence. They responded after the *second* test object if the sequence matched. Thus, we could examine working memory read-out and the animals’ evaluation of the first test object independent of motor activity. As predicted, we found a ramping of gamma bursting in anticipation of working memory read-out. This was coupled with an increase in information specifically about the to-be-tested item and a decrease in beta on channels carrying information. Further, the use of a sequence revealed that gamma and beta played different roles in evaluating different types of matches and non-matches (identity, order) and in corresponding errors. These results lend further support for the hypothesis that discrete oscillatory dynamics underlie maintenance as well as read out of working memory.

## Results

On each trial (Fig. 1) two sample objects were presented sequentially, separated by a 1s delay. Then, after another delay there was a sequence of two test objects, separated by a one second delay. If the identity and order of objects in the test sequence matched that of the sample sequence, the animals were rewarded for releasing a bar. If the test sequence did not match, the monkeys had to wait for one more second and then a matching test sequence appeared requiring a bar release to receive a juice reward (overall performance was 95.5% correct).

**Figure 1.**
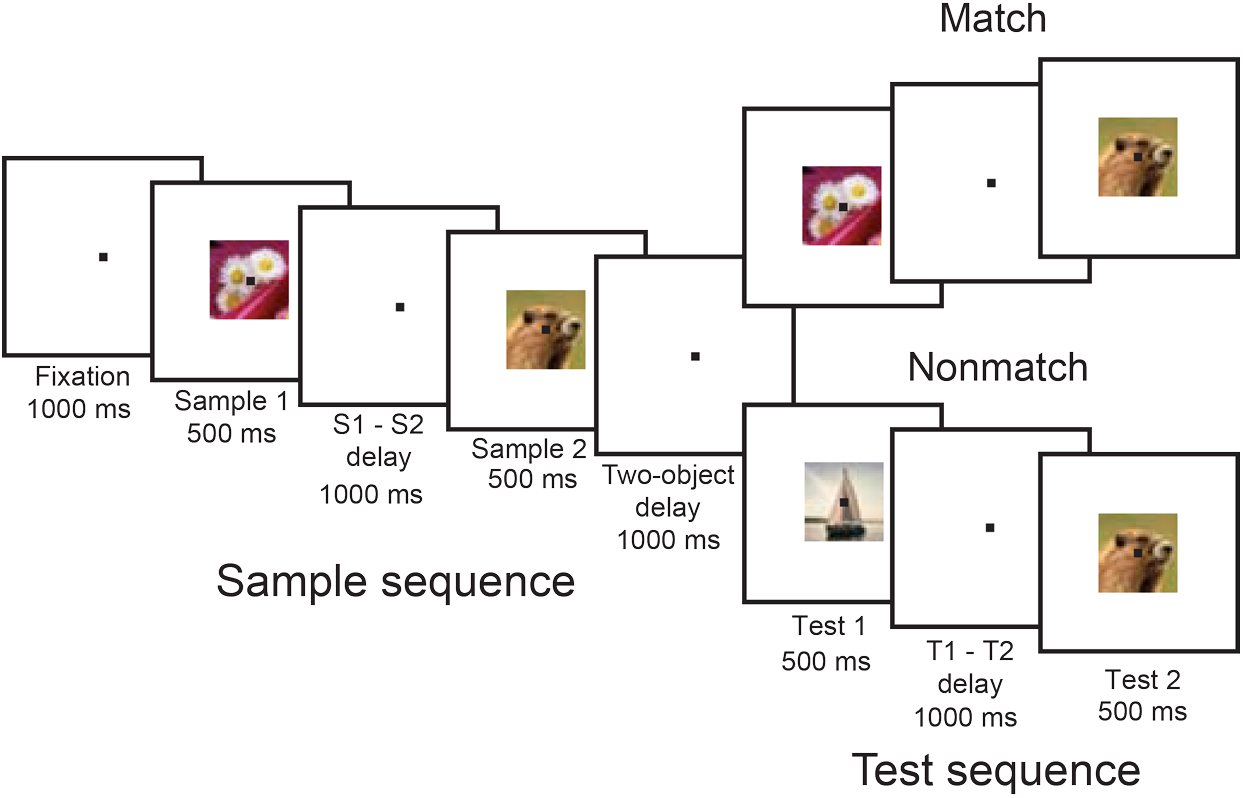
Experimental setup. The animals held a bar and fixated at the center of the screen throughout the task. Two sample objects (S1 and S2) were presented (500 ms), separated by a delay (1000 ms; S1-S2 delay). After another delay (1000 ms; two-object delay) a test sequence of two objects appeared (T1 and T2), separated by a 1000 ms delay (T1-T2 delay). The animals were trained to release the bar after the second test object only if both the identity and the order of the test objects matched the sample sequence. When the sample and test sequences did not match, the animals had to wait for the subsequent (always matching) test sequence to release the bar.

### Information has different relation to gamma and beta bursts

As in prior work^6^, we found that local field potentials (LFPs, n=188 electrodes with at least one spiking neuron) showed bursts of gamma and beta oscillations that were anti-correlated. When gamma was high, beta was low and vice-versa. Virtually all recording sites showed beta oscillations that were elevated during fixation and memory delays and suppressed during object presentations (Fig. 2a). The majority of recording sites (160/188) also exhibited increased gamma bursting when beta oscillations were suppressed (Fig. 2b). These gamma and beta oscillations appeared to be broad band and persistent over time in the trial-averaged data. However, as before^6^, on single trials there were actually brief bursts that were narrow band, and varying in central frequency (Fig. 2c, d). These likely reflected network transitions to high-power states, not noise fluctuations. There were more bursts in the original LFPs than in surrogate data with identical power distribution but randomized phase (Fig. S1; Methods).

**Figure 2.**
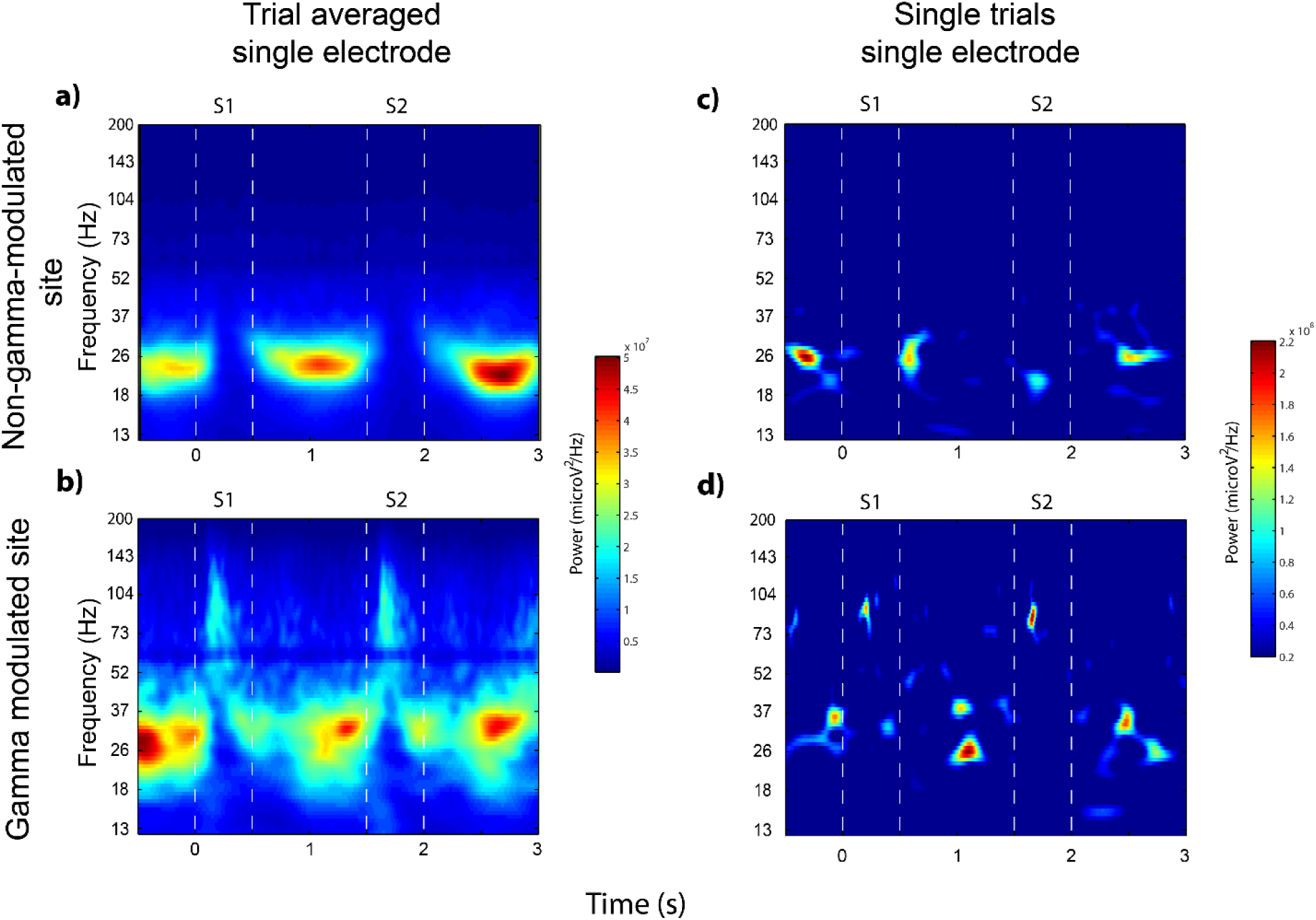
Trial-averaged and single trial spectrograms. Example of trial-averaged spectrograms of a non-gamma-modulated (a) and a gamma-modulated (b) recording site. Displayed are the two sample object presentations, S1 and S2, and the following delays. Single-trial examples originating from the same two recording sites are shown in c) (non-gamma modulated) and d) (gamma-modulated).

Figure 3 shows the gamma and beta burst rates (see Methods) averaged across correct trials with matching test sequences. In particular, Fig. 3a displays the gamma burst rate at the recording sites that exhibited increased gamma bursting during object presentation (“gamma-modulated” - red line) and at the remaining sites (“non-gamma-modulated” - blue line). Not surprisingly, the gamma-modulated sites showed sharp increases in gamma bursting during object presentation while the non-gamma-modulated sites did not. However, in the post-trial period the non-gamma-modulated sites (blue line) showed a higher level of gamma bursting than gamma-modulated sites (red line, Fig. 3a; p<0.0001, two-sided permutation test). Thus, the lack of gamma modulation to objects at non-gamma-modulated sites was not because these sites were incapable of showing gamma bursts.

**Figure 3.**
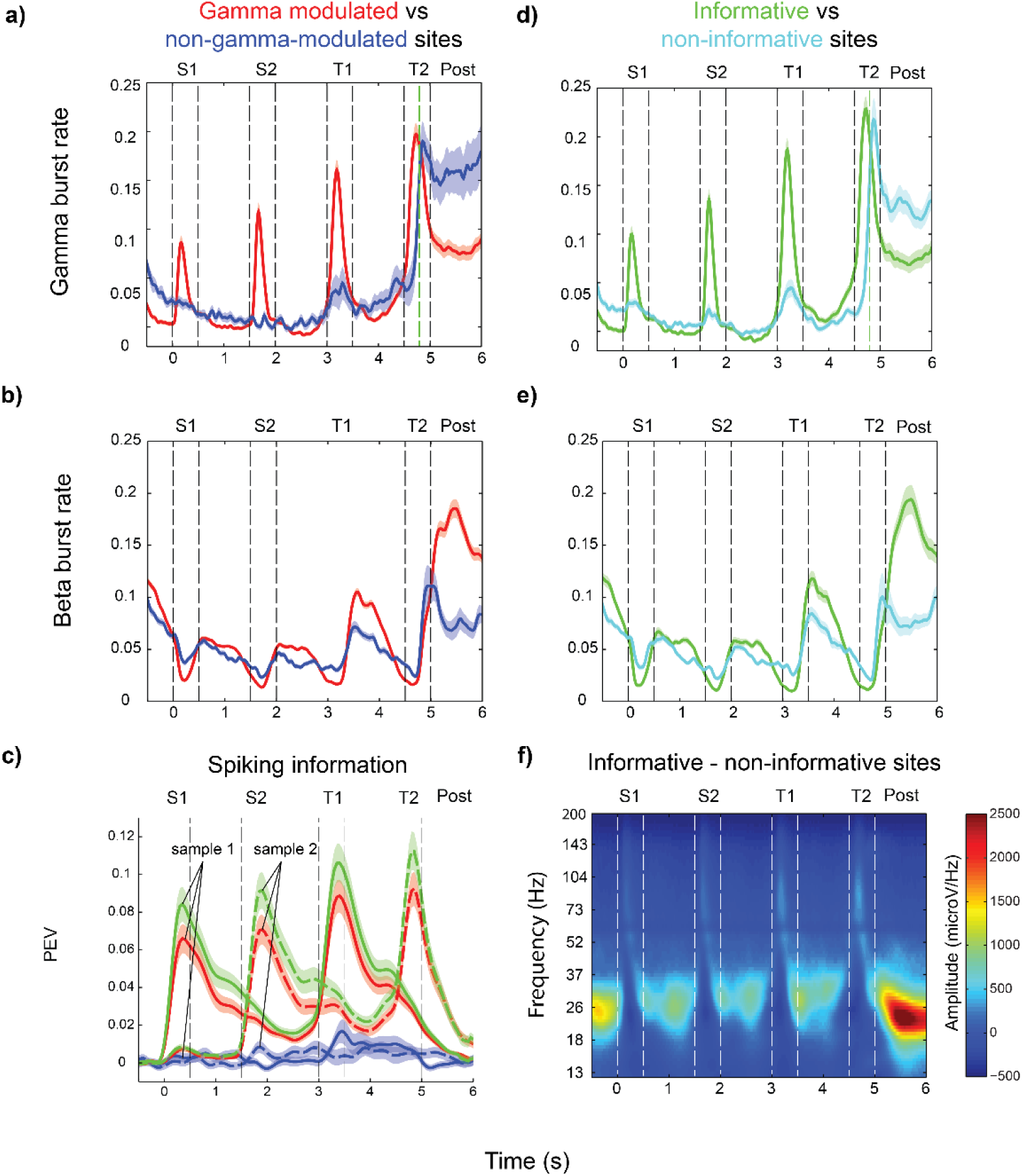
Burst rates and information. All plots show the mean estimates during correct trials with matching test sequences, and 1 s of post-trial (first 1000 ms after T2 offset on correct match trials, vertical green dotted line correspond to the average time of response on these trials) activity. a) Gamma burst rate for gamma modulated (red, n=160) and non-modulated (blue, n=28) sites. b) Same as in a) but for beta burst rates. c) PEV information about the identity of sample 1 (S1, solid lines) and sample 2 (S2, dotted lines) conveyed by firing rates in non-gamma-modulated (blue), gamma-modulated (red) and informative (sites containing at least one unit with significant information (Methods), green) sites. d) Same as in a) but for informative (green, n=130) and non-informative (light blue, n=58) sites. e) Same as b) but for informative and non-informative sites. F) Spectrogram showing average amplitude difference between informative and non-informative sites. T1 and T2 refer to the first and second test object, respectively. Error bars correspond to SEM.

Figure 3b illustrates the average *beta* burst rate at the *gamma*-modulated sites vs non-gamma-modulated sites. The beta burst rate shows the opposite dynamics as the gamma burst rate at gamma-modulated sites (Fig. 3b vs 3a, respectively). Beta bursting was suppressed during object presentation and rebounded during the memory delays, illustrating the push-pull relationship between gamma and beta. This modulation was significantly more pronounced at the gamma-modulated sites (red lines) than the non-gamma modulated sites (blue lines) during both presentations and delay epochs (Fig. 3b, S1: p<0.0001, S1-S2 Delay: p=0.005, S2: p<0.0001, two-object delay: p=0.011, two-sided-permutation test). Recording sites that had the strongest gamma modulation thus also showed the strongest beta modulation (but regulated in opposite direction). The anti-correlation also was present in the post-trial period. For the two groups, the highest post-trial beta burst rate was seen at the gamma modulated sites (Fig 3b; p=0.002, two-sided permutation test), whereas the non-gamma-modulated sites instead had higher gamma burst rates (Fig 3a; p=0.001).

Next, we investigated whether gamma bursts were associated with spiking that carried information about the objects (“informative spiking”), as previously found^6^. For each isolated neuron, we measured information about each of the sample objects by calculating the percentage of explained variance (PEV, see Methods) by object identity. The green line in Fig. 3c shows the PEV averaged across all recording sites with at least one informative neuron (“informative sites”; 130/188). The red line shows the average spike PEV from sites that showed gamma modulation. The PEV from gamma-modulated sites was similar in strength and follows the same dynamic as that of the PEV from the informative sites. This was in contrast to spikes from the non-gamma-modulated sites (blue lines) which showed virtually no information about the sample objects. Thus, object information in spiking was closely associated with gamma modulation by object presentation (p<6e-6, Fisher’s exact test for contingency between gamma-modulated and informative sites).

This is further illustrated in Fig. 3d where the gamma burst rate at sites with informative spiking (green line; see Methods for criteria) vs sites with non-informative spiking (blue line) are plotted. The gamma burst rates from informative vs non-informative spiking sites (Fig. 3d) were virtually identical to the gamma burst rates sorted by whether or not there was gamma modulation (Fig. 3a). While most recorded sites showed gamma-modulation at sample presentation onset, there was a wide distribution in modulation strength. Looking among the population of informative neurons, the amount of information (maximum PEV during sample and delay) correlated positively with stimulus induced gamma (rho=0.42, p<5e-11, Spearman’s rank correlation, n=146) and negatively with induced beta (rho=-0.38, p=5e-6, Spearman’s rank correlation, n=146) during sample presentations. Thus informative sites tended to be the gamma-modulated sites with the strongest modulation of gamma (and suppression of beta).

Figure 3e shows the beta burst rate sorted by whether the sites had informative spiking (green line) vs non-informative spiking (blue line). They were virtually identical to the beta burst rates at gamma-modulated vs non-gamma modulated sites (Fig. 3b). It is important to note that these distinctions were due to whether the spiking was informative or non-informative per se. There were no significant overall differences in average spiking rates between gamma-modulated vs non-gamma-modulated sites (Fig. S2). Thus, as found previously^6^, there seemed to be a tight relationship between gamma modulation by sample object presentation and spiking that was informative about those objects. Gamma however did not appear to be the result of spiking per se. On informative sites gamma and beta bursting were anti-correlated over time (r= -0.40, p<9e-14, n=130, independent samples t-test for sample and delays combined), while on non-informative sites there was no correlation over time (r=0.08, p=0.12, n=58).

During delays, beta bursting increased in informative as well as gamma-modulated sites (Figure 3b, e, green and red lines) relative non-informative and non-gamma-modulated sites, respectively (Fig. 3b, e, dark and light blue lines. Non-informative vs informative sites, S1-S2 delay: p=0.0010, Two-object delay: p=0.04, two-sided permutation tests). We investigated whether this meant that beta, and not gamma bursting, was correlated with information in spiking during the delays. To this end, we selected all neurons with object-selective delay period activity. We correlated their time-dependent PEVs with the temporal profiles of gamma and beta burst rates at the sites on which they were recorded. We found that information on a single-neuron level was positively correlated with gamma across time (r=0.23 in both delays, p<0.00001, t-test, n=146) and uncorrelated with beta bursting (r=-0.07 in first delay, p=0.06, r=0.01 in second delay, p=0.73, n=146) during delays. Within the population of informative neurons most had low PEV values, while a small population was highly informative (Fig. 4). The correlation between gamma burst rates and information over time during delays was driven by this group of highly informative neurons which tended to be strongly correlated with gamma (Fig. 4). Thus, beta was generally elevated at informative sites in memory delays. However, individual neurons only carried information sporadically during the delay^10^, ^11, 12^ (Fig. 5). These informative episodes occurred when gamma bursting on that particular recording site was elevated.

**Figure 4.**
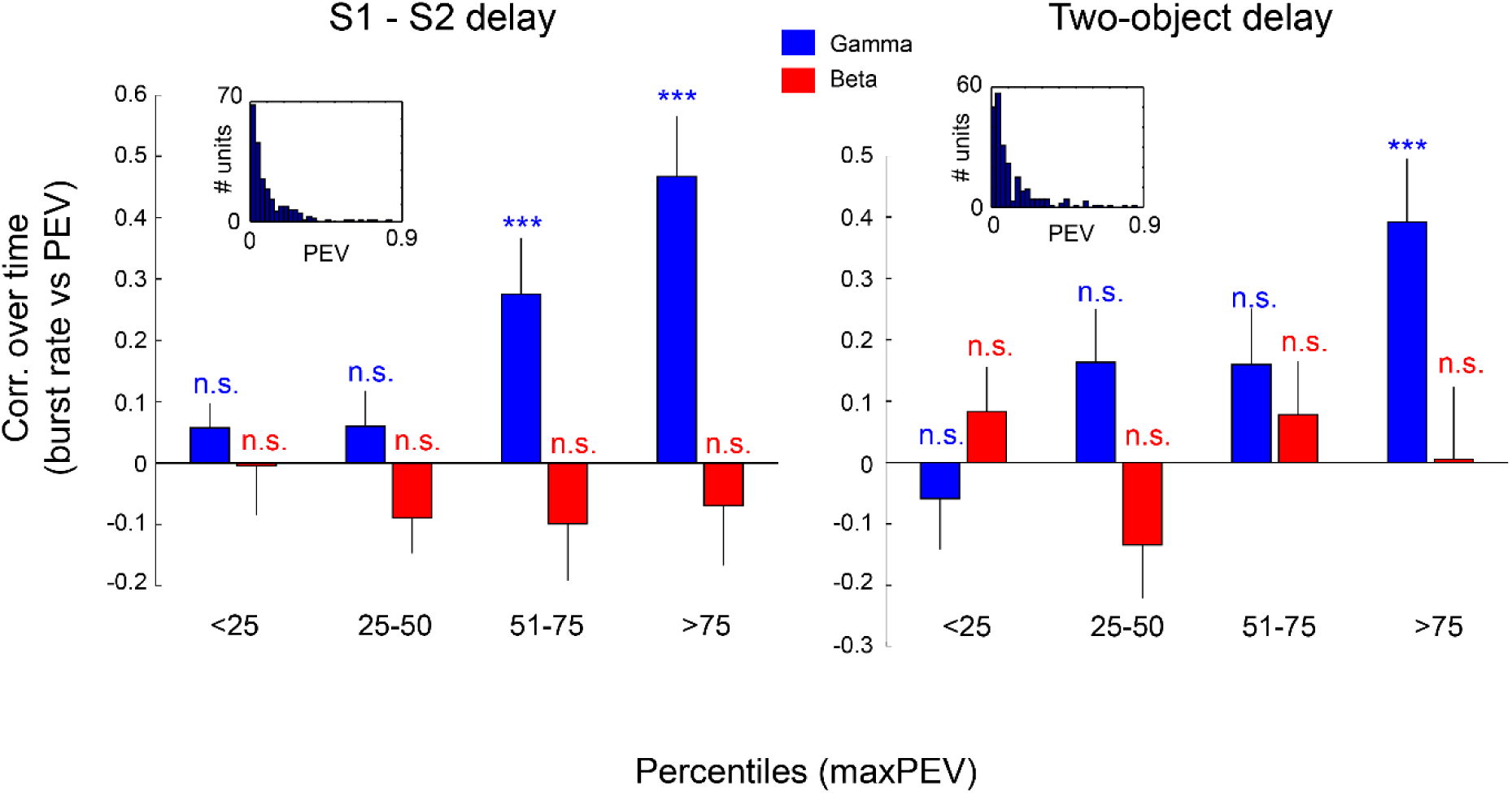
Temporal correlation of PEV information and burst rates. Correlation between temporal profiles of PEV of units with significant delay information (Methods) and gamma (blue) and beta (red) burst rates recorded on the same site. Data are broken down into percentiles based on maximal PEV for each unit. In delay 1 (S1-S2 delay) burst rates were correlated with PEV information about the identity of sample 1 (left), in delay 2 (two-object delay) burst rates were correlated with the sum of PEV information about the identities of both samples 1 and 2 (right). T-test was used with n=36 in each group. In delay 1 only top two quantiles for gamma where significant (p=0.005 and 3.2e-05 respectively), in delay 2 only the top quantile (p=0.001). Error bars correspond to SEM. Inlets show the distribution of peak PEV in each delay for all units with significant information (Methods).

Finally, post-trial gamma bursting (Fig. 3a, d) was higher in the non-informative (and non-modulated) relative to the informative (gamma-modulated) sites (p<0.0001, two-sided permutation test). During the post-trial epoch beta bursting in informative sites reached the highest levels at any point (Fig. 3e, between 5 and 6 sec) while beta in non-informative sites was relatively suppressed. In fact, the most pronounced difference between informative and non-informative sites at any time and frequency was this elevation of beta bursting in informative sites after the end of each trial (Fig. 3f). During this time, information about the second sample object (which was no longer needed) was suppressed (Compare dotted green line at 3 s and 6s in Fig. 3c. Average T2/S2 PEV in the first 1s following T2 offset compared to average S2 PEV following S2 offset, p=4e-7, t=5.3, n=146, t-test).

### Gamma ramp-up reflects information read out

There was a ramp-up of gamma bursting just before presentation of the first (T1; p<0.0001, Fig. 3a, d) and second test object (T2, p<0.0001, informative sites, n=130, two-sided permutation test on informative sites, n=130, for all comparisons here, ramp-up was evaluated comparing statistics between two 300 ms epochs – the last 300 ms of a delay and the preceding 300 ms). This was especially strong for gamma from informative sites (Fig. 3d, green line). Similarly, while beta tended to be elevated at informative sites in comparison to non-informative sites during delays (Fig. 3e), it was relatively suppressed just prior to tests (T1; p<0.0001, T2; p<0.0001, last 300 ms of delays) but not samples (S2; p=0.12). There was a corresponding ramp-up of object information in spiking (PEV) before tests. Objects were tested one by one, in a sequence. Information only ramped up for the sample item that was about to be tested, i.e. information about the identity of the first sample object ramped up before the first test object, and information about the second sample object ramped up before the second test object (Fig. 3c, PEV about S1 before T1, p=3e-5, t=4.3. PEV about S2 before T2, p=3e-5, t=4.2, n=146). For objects in the sequence that were not relevant in the upcoming test, there was instead a non-significant decreasing trend (PEV about S2 before T1, p=0.08, t=-1.8; PEV about S1 before T2, p=0.10, t=-1.7, n=146). To investigate this further, we sorted neurons by when in the delays they carried information (Fig. 5). If a given object was not the one being tested next (e.g., the *second* sample object in the delay before the *first* test object), neural information about that object was evenly spread over the delay (Fig. 5 a, c-d). If the object was the next one to be tested (e.g., the first sample object before the first test object), neural information about that object tended to “pile up” at the end of the delay before the test (Fig. 5 b, e).

**Figure 5.**
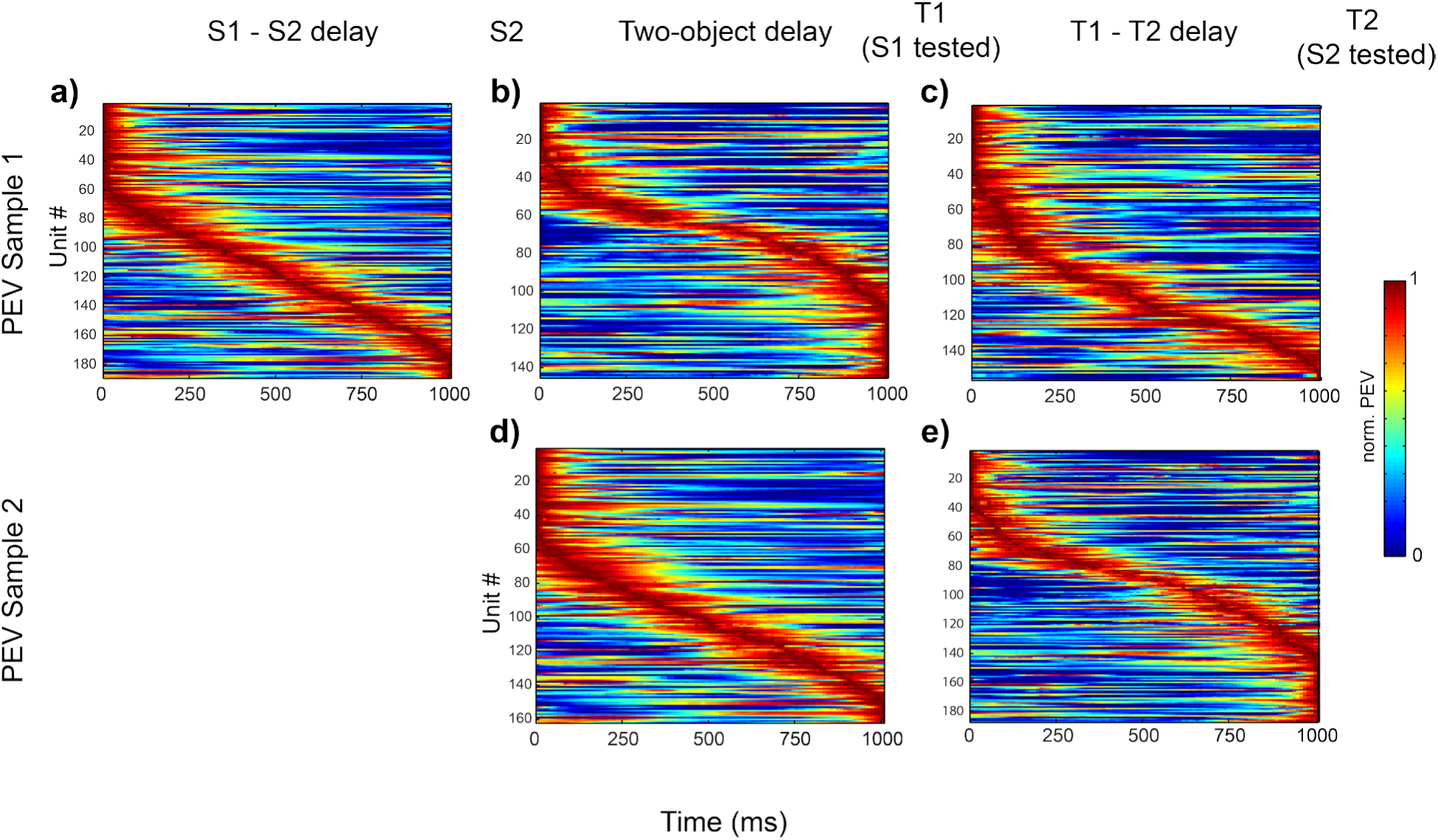
PEV information during different types of delays. Data from trials with matching test sequences. Plotted is the normalized PEV information about sample 1 (S1; top panels) or sample 2 (S2; bottom panels) during S1-S2 delay (left column), two-object delay (middle column) and T1-T2 delay (right column) for all units with significant information (p<0.01; ANOVA, tested in each delay and for each sample independently). Units are sorted based on the timing of peak PEV in each delay. Following two-object delay, S1 is tested. Following T1-T2 delay, S2 is tested.

We interpreted the ramp-up of gamma bursting as read-out of working memory in anticipation of the decision about matching vs non-matching type of test stimuli. The ramp-up did not account for prediction of a presentation of any object. It only occurred before the test objects. There was no ramping of gamma bursts (rather a non-significant decrease prior S2; p=0.10, n=130) or spiking PEV before presentation of the first or second sample objects (Fig. 3c and 3d), even though both events were predictable. In the case of the ramp-up prior to the first test object, we can also rule out motor planning because no behavioral response could be determined until the second test object. Finally, the motor response was always the same (a bar release). Thus, motor activity could not explain that *information* only about the to-be-tested object increased.

In sum, these results suggest that gamma and beta bursting have a push-pull type of relationship. When gamma bursting is high, beta bursting tends to be low. It also illustrates that modulation of gamma bursting is closely associated with informative spiking, over time as well as across sites. Next, we show that gamma and beta also differed in how they related to the match/non-match status of the test objects.

### Gamma and beta shows different responses to matches and non-matches

Our task required matching a sequence of two test objects to a sequence of two sample objects. This allowed us to examine neural activity associated with different types of non-matches (object order vs identity). We focused on the first test object and the following delay since there was no behavioral response during these epochs.

If the first test object did not match either of the sample objects, it was termed an “object non-match”. If the first test object matched the *second* sample object, we called it an “order non-match”. We found that gamma bursting during test object presentation distinguished between a match and different types of non-matches (Fig. 6a). During presentation of the first test object, the gamma burst rate was lowest for a match, highest for an object non-match and intermediate for order non-match (horizontal black lines denote intervals when burst rates for object non-match and match were significantly different, p<0.05; cluster based statistics).

**Figure 6.**
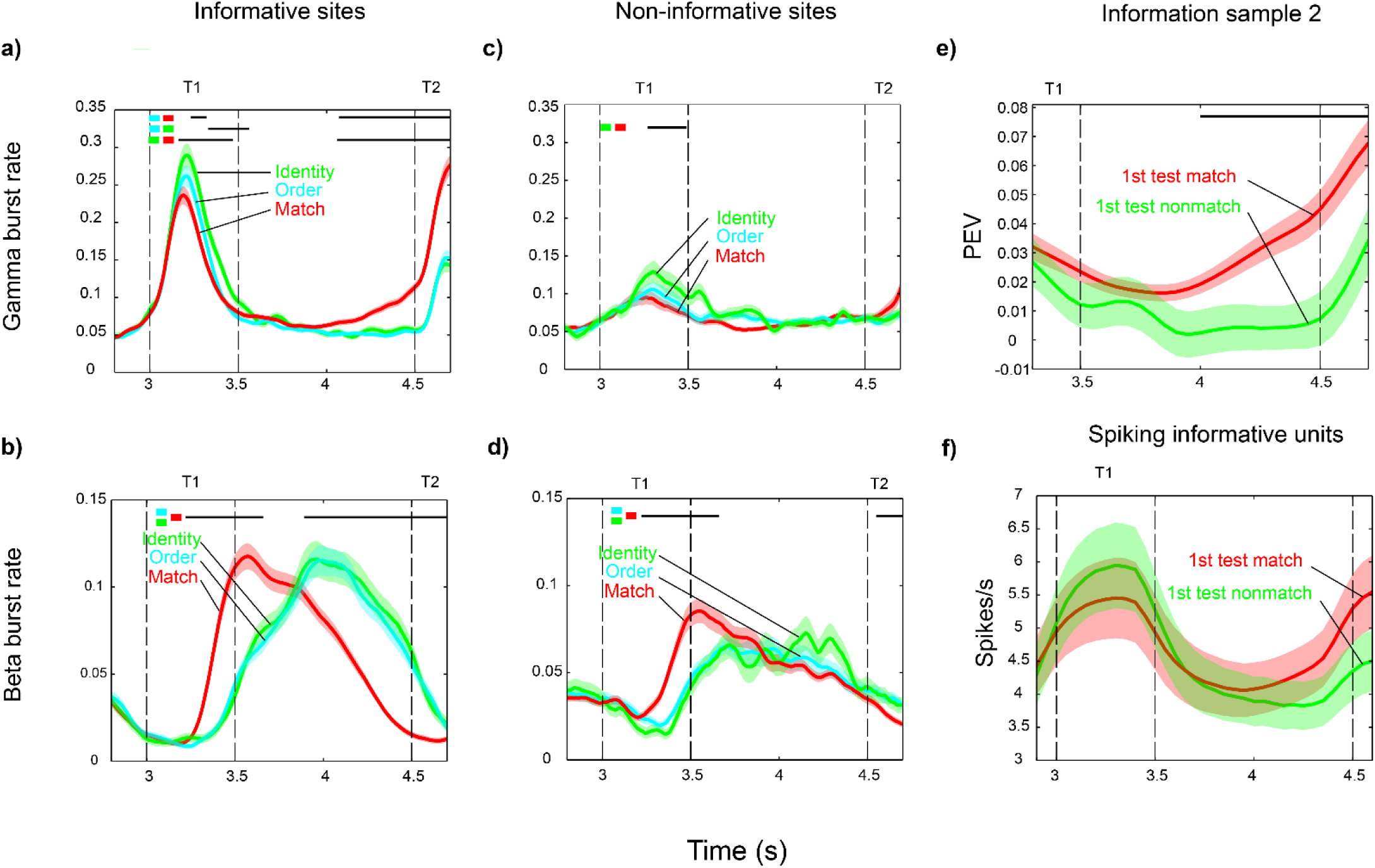
Burst activity around the first test. Plots show burst rates during presentation of the first test object (T1) and the following delay for correct trials. a) Gamma burst rate for matching (red) and non-matching sequences (object identity violation, green; order violation, blue) in sites with at least one unit carrying significant information about the identity of sample 1 or 2 (n=130). Black horizontal lines denote intervals with significant differences between burst rates in conditions indicated by the corresponding colored squares (p<0.05; cluster based statistics with permutation test). b) Same as a) but for beta burst rate. c) same as a) but for sites with no informative units (n=58). d) Same as b) but for sites with no informative units. e) PEV information about the identity of sample 2 (tested after the delay at T2) for units with significant information during delay (n=146) in two groups of trials: with matching (red) vs. non-matching (green) pair of objects, S1 and T1 (at first test). Black horizontal line marks an interval where the PEV means in the two groups of trials are significantly different (p=0.001; one-sided cluster based permutation test). f) Average firing rates at informative sites (n=189) for first test matching (red) and non-matching (green) conditions (analogously to e)). Error bars correspond to SEM.

By contrast, beta distinguished between matches and non-matches in the timing of the beta bursts in the delay following the first test object (Fig. 6b, black line denotes significant differences between match relative both object non-match conditions, p<0.05; cluster based statistics). However, it did not distinguish between different types of non-matches. There was no significant difference between order and identity non-matches. The differences in gamma bursting between conditions were short-lived and disappeared shortly after test object presentation. The differences in beta were of longer latency as well as duration, and bridged the one-second delay to the second test object.

Gamma burst rate also ramped up as the presentation of the second test object approached, but only if the first test object was a match (and thus further read-out was needed, Fig. 6a, red line). If the first test object was either type of non-match (blue and cyan lines), there was no gamma ramp-up (Fig. 6a). Presumably the animal could have already rendered its non-match decision after the first test object was a non-match. Indeed, the corresponding ramping up of information in spiking about the second sample in informative neurons (Fig. 3c) occurred only if the first test object was a match (Fig. 6e; p=0.001, cluster based statistics with permutation test). As above, all these effects were stronger at informative sites (Fig. 6a, b) than at non-informative sites (Fig 6c, d). The gamma ramp-up before the second test object in match trials and the corresponding increase in beta bursting in non-match trials were only seen on informative sites. Non-informative sites showed no significant differences (Fig. 6c, d). Spike rates on informative sites showed similar tendencies as gamma bursting, but with no significant differences between match and non-match conditions (Fig. 6f).

### Gamma and beta reflects different types of errors

We examined trials in which the monkeys made errors in match/non-match judgments (i.e., the wrong behavioral response after the second test object). For this analysis, we focused on informative sites because they showed the most robust effects (see above). We combined both types of non-match trials (object and order) to obtain enough incorrect trials for statistical analysis.

Figure 7 shows the average gamma and beta burst rates on error trials relative the correct trials. Plotted is the gamma (Fig. 7a, b) and beta (Fig. 7c, d) burst rates from correctly performed match trials (red) and correctly performed non-match trials (blue). Plotted in black (Fig. 7a, c) are the burst rates on trials in which the first test object (and thus the sequence) was a non-match, but the monkey mistakenly responded “match” to the second test stimulus. In Fig. 7b and d the black curves denote the burst rates on incorrect trials in which the test sequence matched the sample sequence, but the animal mistakenly classified it as a non-match (by not responding). Below each figure are horizontal lines indicating intervals with statistically significant differences (p<0.05; cluster based statistics) between the correct match and correct non-match trials. Within each interval, we then calculated the average number of gamma or beta bursts (per trial and per electrode) for incorrect and correct trials. The second gamma burst rate interval was divided into two, as it overlapped both the delay and presentation of the second test object.

**Figure 7.**
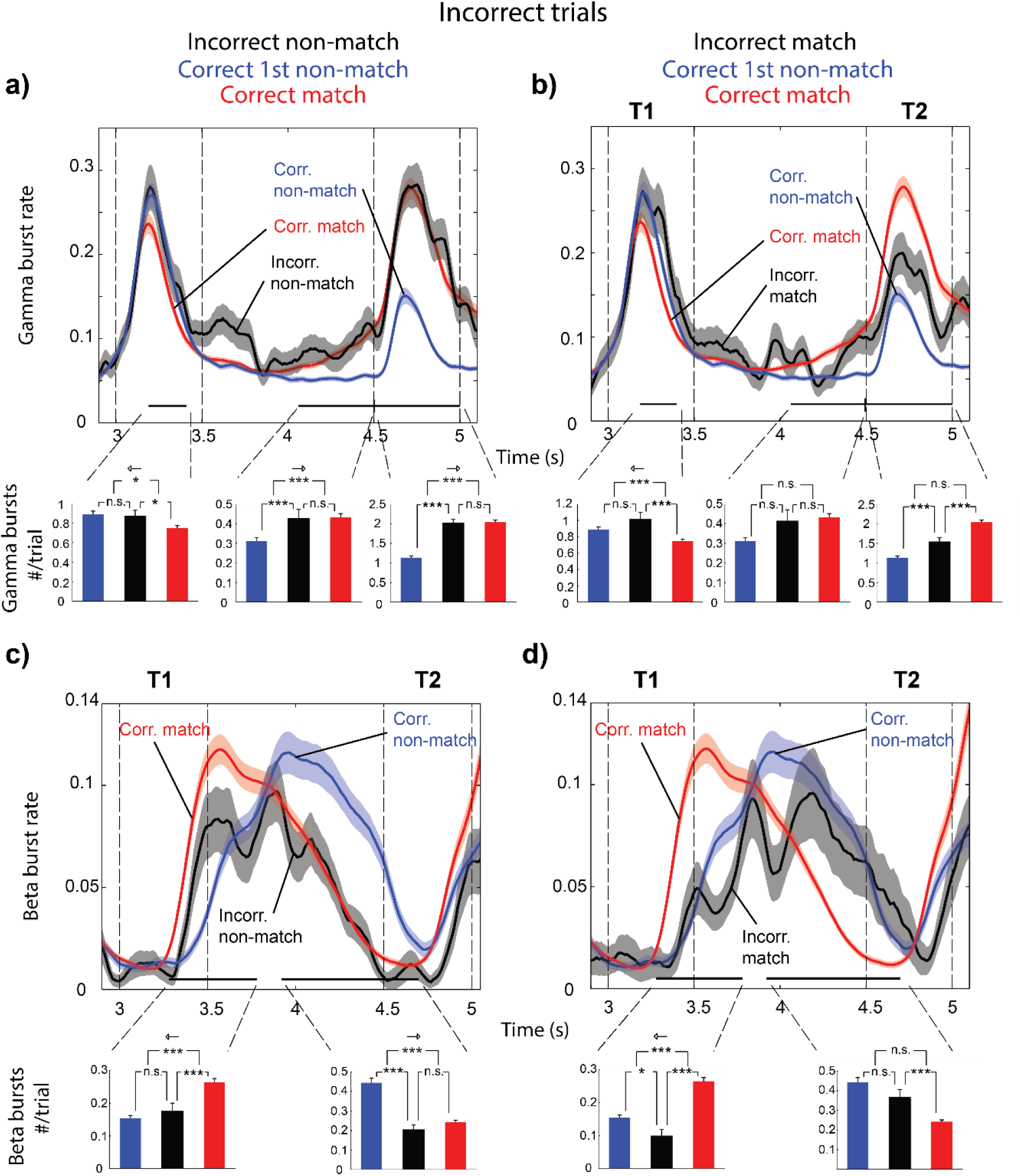
Bursts dynamics for incorrect trials. Bursting dynamics during and following the presentation of the first test object, T1, in informative sites (n=130). The corresponding burst rates for the correct matching (red) and correct first test non-matching, i.e. T1≠S1 (blue) conditions are given as reference. Black curves illustrate burst rates for incorrect trials in the matching (right panel column) and first test non-matching (left panel column) conditions. Black horizontal lines at the bottom of each panel denote intervals when the burst rates in the correct matching and correct non-matching conditions are significantly different (p<0.05; cluster based permutation test). Bar plots represent the average number of bursts per trial and recording site for the three conditions in the corresponding panel within the aforementioned intervals. Planned pair-wise comparisons between these mean statistics (the incorrect condition was compared to the two correct conditions) were conducted using two-sided permutation tests. In addition we tested the hypothesis that burst counts on incorrect trials were equidistant (Methods) to the two correct conditions (arrow indicate which condition the error trials were more similar to for significant effects). a, b) show gamma burst rates, and c, d) show beta burst rates. For correct trials, n=130 (number of electrodes) whereas for the incorrect matching trials, n=117 (as some sessions were devoid of error trials of this kind) and for the incorrect non-matching trials, n=118. Error bars correspond to SEM.

First, we consider errors when the first test object was a non-match (Fig. 7a, c). During the first test object, the gamma burst rate on these trials (black curve) overlaps with the gamma burst rate from correct non-match trials (blue curve). The number of gamma bursts was significantly different from that in correct match trials but not from that in correct non-match trials (Fig. 7a, bottom) during the first test. The same was true for beta bursting (Fig. 7c). Thus, it seemed that in trials in which the monkeys mistakenly responded “match” to a non-matching sequence, the gamma and beta burst rates during the presentation of the first, non-matching test object followed the “correct” trajectory (i.e., as if the first test object was a non-match). Instead, the error seemed to arise in the delay after the first test object. Interestingly, in that delay period there was a ramp-up of gamma burst rate on incorrectly performed non-match trials (black line) which closely followed the gamma burst rate on correctly performed match trials (red line). During the last part of the delay leading up to the second test object, the average number of gamma bursts on incorrectly performed non-match trials (Fig. 7a, bottom, black bar) was not significantly different from correctly performed match trials (red bar), but was significantly different from correctly performed non-match trials. This was reflected also in the beta bursting, which in the latter half of the delay was significantly suppressed relative correct non-match trials. The average number of beta bursts was significantly lower than the number in correct non-match trials, but not significantly different from correct match trials (Fig. 7c, bottom). In sum, if the first test stimulus was a non-match, the gamma and beta burst rates followed the average trajectory observed for correct identification. The error in responding “match” seemed to occur in the second half of the delay as the gamma and beta burst rates become more similar to the “match-like” profiles.

Next we looked at trials with matching test sequences, where the monkey failed to respond, as if the sequence was a non-match (Fig. 7b, d). In this case, the gamma burst rate to the first (matching) test object (black line) virtually overlapped with that in correct trials when the first test object was a non-match (blue line). The number of gamma bursts during the first test object (black bar) was not significantly different from that on correctly performed non-match trials (blue bar), even though the first test object was actually a match (Fig. 7b). The number of beta bursts was likewise significantly lower than that of correct match trials (Fig. 7d). In the second half of the delay, leading up to the second test, beta bursting was not statistically different from that in correct non-match trials and significantly higher than that in correct match trials. There was no significant difference in gamma bursting between error (black line and bar) and the correct match (red line, bar) nor non-match trials (blue line, bar). During the second test object, however, the error gamma burst rate was intermediate to, and significantly different from correctly performed match (red bar) and correctly performed non-match trials (blue) bar.

In sum, on trials in which the first test object was a non-match, the gamma and beta burst rates initially followed the trajectories corresponding to the trials with the correct non-match identification. The error of recognizing the sequence as a match crept in later in the run-up to, and during presentation of the second test object. On trials in which there was a matching sequence but the animals mistook it for a non-matching sequence, the gamma burst rate on error trials followed “wrong” dynamics from the start, during presentation of the first test object. Likewise, the beta burst rate during the delay was as if the first test object was a non-match, even though it was a match.

Finally, we wanted to rule out that differences in burst rates between conditions was due base-line shifts in power. For all intervals in Fig. 7 in which we tested for significant differences in burst number between the conditions (correct match, incorrect match and both sets of error trials), we also tested for differences in average power during bursts (Table. S1; Methods). There were no significant differences in power during beta bursts between any conditions. For gamma there was only significant differences at the second test, between conditions where the animals did a motor response for one condition but not the other. This demonstrated that changes in burst rates were not the result of thresholding of signals with different tonic means. Instead, it suggests that the behavioral correlates of matching and non-matching test objects were changes in the rate of burst occurrences.

## Discussion

We analyzed LFPs and spiking activity during a working memory task^8^, ^9^. The task structure gave us an opportunity to examine activity as monkeys read out an object sequence from working memory and compare it to a sequence of test objects. We found brief gamma and beta bursts that seemed to have different functions, confirming and extending previous results^6^. Gamma bursts were temporally and spatially linked with the expression of sensory information in spiking during encoding and delays. Beta bursts were associated with suppression of gamma and suppression of object information in spiking. Gamma and beta bursting were anti-correlated over time, but only at recording sites where spiking carried information about the to-be-remembered objects (informative sites).

The beta and gamma interplay suggests a potential mechanism for controlling working memory^6^, ^7^. The balance between beta and gamma would control the level of gamma bursting and hence, the expression of sensory information in spiking linked to gamma. The results suggest that beta/gamma balance is under top-down control. It was, along with information, modulated by task demands absent of sensory stimuli. The biggest difference between informative and non-informative sites was after the behavioral response, in the post-trial epoch. At informative sites, beta was particularly high and gamma low. At this point, the memory content is no longer relevant. Thus, the shift to activity dominated by beta may clear out working memory by suppressing gamma. Indeed, at the same time, spiking information about the last object held in working memory (the second sample/test object) decreased dramatically, as if suppressed.

Beta and gamma activity during working memory read-out and match/non-match decisions were consistent with their suggested roles in regulating working memory. The rate of gamma bursts ramped up, and beta rates decreased, at the end of memory delay in anticipation of the comparison of the memories to the forthcoming test objects. This was accompanied by an increase in end-of-delay spiking that is often seen in WM tasks^13^, ^14^, ^15^. Here, we observed it in absence of, and unrelated to, any forthcoming motor response, as did Hussar and Pasternak (2010). Further, we found the ramp-up in spiking carried the specific object information needed for the immediately forthcoming decision (e.g., first sample object information for comparison to the first test object, etc). The gamma/informative spiking ramp-up and beta ramp-down did not occur in anticipation of just any expected event. Sample object presentation was also predictable, but no ramp-up was present. Thus, gamma ramp-up coincides with working memory read-out, but not encoding. It also did not occur before the second test object, if the first test object was a non-match. This rendered the whole sequence a non-match and the second test object was no longer relevant. Thus, there was no gamma or informative spiking ramp-up about this object. Instead, there was an increase in beta at sites with information.

These dynamics continued to play out in the comparison between the memories and the test objects. Gamma bursting was highest for identity non-matches, second highest for order non-matches and lowest for matches. This reflects their relative level of “non-matching”. It mirrors observations of changes in average spike rate in prefrontal cortex^4^,^16^. The changes in gamma were then followed by changes in beta. When the first test was a match, beta was elevated immediately after its offset, perhaps clearing the memory of the first object so the second one could subsequently be read out. When it was a non-match, the beta increase came later, only at informative sites and just before the second test object, as if preventing read-out of the now irrelevant second object.

Deviations from these match and non-match gamma/beta dynamics predicted behavioral errors. When the first test object was a non-match, initial gamma bursting reflected its non-match status, whether or not animals made an error. The “match error” instead crept into both gamma and beta bursting later, in the delay between the two test objects. Then gamma and beta bursting reached levels similar to when animals correctly identified a match. There was a gamma increase and a beta suppression. It was as if animals expected to make decision about the second test object. However, that was only necessary if they thought the first test was a match, not the non-match that it was. When, instead, the first test object was a match and animals subsequently responded non-match, the error was instead immediate. The gamma (and subsequently) beta burst rate to that object was similar to non-matching test objects, instead of the match that it was.

Cortical gamma has long been seen as a correlate of sensory processing^17^ but the role of beta has been more elusive^18-23^. Beta has been suggested as an inhibitory rhythm^18^, ^19^, to be involved in motor maintenance^20^, post-movement rebound^21^, ^22^ or a mechanism to preserve status quo^23^. The interplay between oscillations and spiking seems congruent with an inhibitory role of beta. Increases in beta were correlated with suppression of gamma and informative spiking. Beta was also elevated post-trial, when information needed to be cleared out. This could explain why motor beta is most pronounced after a completed movement^21^, ^22^, when the movement plan should be forgotten. In the human ventral stream, different patterns of gamma were induced by different visual stimulus categories, while beta is globally reduced^24^. It is therefore possible that higher-order cortex added volitional control onto mechanisms similar to that found visual cortex. It should be noted that the beta observed in this study was in the high beta (β2) range, and beta oscillations in the β1 frequency band might have other behavioral correlates^25-27^.

Sustained spiking has often been seen as the neural correlate of working memory^1-4^. It has been modelled by attractor networks with persistent activity^5^. The activity in such networks is by definition stable to perturbations. Here the observed dynamics were not sustained, but occurred in brief bursts. Dynamically speaking, brief bouts of gamma and informative spiking, with interleaved periods of silence might be a way to combine the robustness of attractor-like activity with more flexible computations^28-31^. If gamma bursts correspond to periods of short-lived attractors^7^, the periods of silence between them might be opportunities for the network to evolve and weave in new information. Time-varying signals in working memory delay activity appears to be a hallmark of prefrontal dynamics^4, 9, 10, 12, 31-33^. We suggest that fast transitions between brief high-power events in gamma and beta allow for the flexible coordination of multiple items held in working memory.

## Methods

### Behavioral task

The task was structured such that the animals had to compare one working memory item at a time to test stimuli. Each trial (Fig. 1) consisted of an encoding phase in which two objects were presented sequentially, separated by a 1s delay. The second sample object was followed by another delay and then a sequential test phase. The identity as well as the order of the items in the test sequence had to match that of the to-be remembered sequence. The monkeys reported a matching test sequence by releasing a bar. If the test sequence did not match, the monkeys had to wait for a second, matching test sequence before releasing the bar and receiving a juice reward (overall performance was 95.5% correct). Throughout the whole trial the animals had to fixate on a dot in the center of the screen and all items were presented in the same location, at the fixation dot (Figure 1). Thus, item information was not confounded with location information and planned saccades not any part of the task.

### Data collection

For each recording, a new set of acute electrodes (up to 8 simultaneously) were lowered through a grid. The LFPs were recorded at a sampling rate of 1 kHz. For details please see Warden & Miller 2007^8^ as well as Warden and Miller 2010^9^, in which the original data was recorded.

### Signal processing

At first, all electrodes without any isolatable neurons were removed. Then, a notch filter with constant phase across a session was applied to remove 60-Hz line noise and its second harmonic. Two methods for the LFP spectral estimation were employed: Morlet wavelet analysis^34^ and multi-taper approach with a family of orthogonal tapers produced by Slepian functions^35^, ^36^. They yielded very similar results in terms of qualitative time-frequency content. They also led to comparable burst extraction outcomes. For all the presented spectrograms (except Fig. 2 where wavelets were used) and for burst analysis the multi-taper approach was adopted with frequency-dependent window lengths corresponding to six to eight oscillatory cycles and frequency smoothing corresponding to 0.2-0.3 of the central frequency, *f*_0_, i.e. *f*_0_±0.2*f*_0_, where *f*_0_ were sampled with the resolution of 1 Hz (this configuration implies that two to three tapers were used). The spectrograms were estimated with the temporal resolution of 1 ms.

### Burst extraction and detailed estimation of power inside bursts

The bursts were calculated similarly as in the previous study^6^ with the only difference in estimating the reference mean and standard deviation of spectral band power. Here, the statistics were obtained over the ten-trial-long period (the last nine plus the current trial) to minimize the potential effects of removing true effects in trial to trial differences in power between conditions.

The first step of the oscillatory burst identification consisted in extracting a temporal profile of the LFP spectral content within a frequency band of interest. We used single-trial spectrograms, obtained with multi-taper approach, to calculate smooth estimate of time-varying band power. Oscillatory bursts were recognized as epochs during individual trials when the respective measure of instantaneous spectral power exceeded the threshold set as two SDs above the mean of the respective band power over the ten-trial-long reference period, providing that they lasted at least three oscillatory cycles (for the mean frequency of the band of interest). To obtain a more accurate estimate of burst duration, the time-frequency representation of the signal was extracted in the spectro-temporal neighborhood of each burst using the aforementioned multi-taper method, and two-dimensional Gaussian function was fitted to the resulting local time-frequency map. The burst length was then defined as a time subinterval where the band average instantaneous power was higher than half of the local maximum (half-power point) estimated using the Gaussian fit. The frequency coordinate of the peak of the Gaussian fit was recognized as the central burst frequency. For each burst the spectro-temporal power average was calculated and normalized with reference to the session power spectral average within a narrow band around the central frequency of a given burst.

Finally, based on burst intervals extracted from each trial for the beta band (20-35 Hz) and two gamma sub-band oscillations (55-90 and 80-120 Hz), we defined for each band a trial-collective measure, called a burst rate, as the proportion of trials where a given electrode displayed burst-like oscillatory dynamics around the time point of interest sliding over the trial length. In other words, a burst rate corresponds to the time-varying likelihood of a burst occurrence on a given electrode at a specific time point in the trial. Burst rates were estimated for beta and gamma sub-bands. For all figures and statistics of gamma burst rate we used the summed burst rates of the two sub-bands.

### Testing the bursty nature of data

We tested that the data contained more high power bursts than could be expected from random fluctuations around the same spectral power content. In the first step we did a Fourier decomposition of the data, and then randomized the phases of the components^22^. We then applied an inverse Fourier transform to obtain surrogate signal with preserved distribution of power but with random temporal relationships. To this signal we then applied the same signal processing and burst extracting procedure as to the original signal. This was repeated 250 times to obtain a distribution of estimated burst rates in surrogate data relative original data (Fig. S1; estimated burst rate was always higher in the original data).

### Selection of informative cells/sites

The instantaneous firing rates were estimated for each neuron by convolving spike trains with a Gaussian kernel (30-50 ms wide). The bias-corrected PEV^37^, ω^2^, was then estimated from firing rates with the resolution of 1 ms across trials with different stimulus dependent conditions. As a result, PEV allowed for the quantification of information associated with the modulation of firing rates (variance) of individual neurons depending on the stimulus condition. This way, for example, we could estimate the amount of variance-based information carried by individual neurons about the identity of the presented object. The bias-correction minimizes the problem of non-zero mean PEV for small sample sizes.

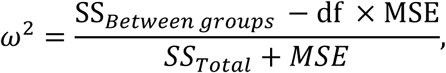
where *MSE* is the mean squared error, df the degrees of freedom, *SS_Total_* the total variance (across all trials) and *SS_Between groups_* the variance between groups of trials formed wrt. stimulus condition of interest. We used one-way and multi-way ANOVA for the condition of interest to recognize informative neurons, with very similar results. A neuron was defined as informative if the ANOVA analysis provided statistically significant evidence (p<0.01, Bonferroni corrected for testing at multiple time-points) for the rejection of null hypothesis, thus yielding relevant group dependent effects (*ω^2^*) at any time point during presentation and delay periods.

### Statistical methods

The majority of tests performed (all burst rate comparisons and PEV comparisons) in this study were nonparametric due to insufficient evidence for model data distributions. To address multiple comparisons problem we employed permutation, Friedman’s and Wilcoxon’s signed-rank tests where appropriate. We also performed Pearson’s rank correlation or students T-test for non-zero mean for correlations between burst rates and PEVs. For details see below.

### Correlation

We also estimated the correlations between the measures of time-varying spectral band content, burst rate statistics and PEV profiles over time in WM delays. These measures are by definition estimated over a set of trials (collective measures) and we used trial averaged signals on individual recording sites. For example, gamma burst rate was correlated with the beta burst rate and spike PEV over time, on the same electrode.

In addition, we correlated information with induced gamma and beta bursting. To this end we calculated the average burst rate across all presentations (500 ms) divided by the average of all preceding epochs of fixation (500 ms). For PEV information we estimated the maximum PEV value during the presentation and the following delay. For each neuron we thus obtained one data point. Next, we correlated the resulting data points across the population. To mitigate the biased effect of non-uniform distribution of PEVs (a large number of close-to-zero values and a low number of high values), we resorted to Spearman’s rank correlation.

Finally, some attention should be given to the way we report correlations between the measures of time-varying spectral band content, burst rate statistics and PEV profiles. The correlation analyses were performed on individual electrodes and only the summary statistics (mean and SE) were presented.

### Error trial analysis

In order to investigate whether incorrect trials exhibited similar burst characteristics as in matching or non-matching correct trials in different epochs during the test period (T1, T2 and the delay between T1 and T2), we performed the following analysis. First, we defined intervals of potential interest based on the statistical comparison of temporal profiles of burst rates in correct non-matching vs correct matching trials using a permutation test on the largest cluster based statistics^38^ at the significance level of 0.05. This approach allowed for increasing the test sensitivity based on the assumption of temporal continuity of the data, thereby avoiding a massive multiple comparison problem and resulting in continuous intervals. These intervals were calculated separately for gamma and beta burst rates. Second, for each of the resulting intervals we extracted various burst characteristics, i.e. average number of burst occurrences, their average duration and their average spectral power (over the duration and, respectively, beta or broad-band gamma frequency range). Finally, these burst statistics were compared within the intervals using a nonparametric permutation test for exchangeability of condition labels. We also tested if error trials behaved more similar to either of the correct conditions: Within each interval of interest we calculated two statistics: i) from the difference between burst rates in correct non-match minus incorrect conditions and ii) difference between burst rates in incorrect minus correct match conditions. Finally, we tested the null hypothesis that the two means were the same.

The number of error trials was low, i.e. on average there were 4.7 non-matching and 4.1 matching incorrect trials per session.

